# Eusocial evolution without a nest: kin structure of social aphids forming open colonies on bamboo

**DOI:** 10.1101/2022.10.23.513187

**Authors:** Keigo Uematsu, Man-Miao Yang, William Amos, William A. Foster

**Author notes:** Author for correspondence: Keigo Uematsu. Department of Biology, Keio University, Hiyoshi, Kanagawa, Japan (Keigo Uematsu).

## Abstract

Living in nests is an almost universal feature of eusocial animals. In some aphids, however, sterile soldier castes have evolved in open colonies without a nest. To clarify the factors promoting the evolution of eusociality in these colonies, we used newly developed microsatellite markers to compare the kin structure of the open colonies of two aphid species on bamboo: the non-eusocial colonies of *Astegopteryx bambucifoliae* and the eusocial colonies of *Pseudoregma alexanderi* on *Dendrocalamus latiflorus*.

Our samples, from over 1,000 hectares, contained 99 clones of *A. bambucifoliae* and 19 of *P. alexanderi*. Clonal mixing occurred in both species: average pairwise relatedness within a colony was 0.54 in *A. bambucifoliae* and 0.71 in *P. alexanderi*. Each clone of *A. bambucifoliae* occurred in a unique location, whereas those of *P. alexander*i occurred in multiple locations and more than 90% of individuals came from just four clones. There was significant genetic variation among different colonies in the same clump (stem-cluster) in *A. bambucifoliae* but not in *P. alexanderi*, indicating that *P. alexanderi* colonies in a single clump are genetically homogenized, functioning as a large colony. In *P. alexanderi*, the proportion of sterile soldiers to normal first-instar nymphs was significantly different across the four clones.

Our results indicate that the lack of input of migrants from the primary host and feeding on a large, stable host plant are important ecological factors that might favour the evolution of eusociality, enabling the production of genetically homogenised, large, and long-lived colonies. After eusociality evolves on the secondary host, the optimal strategy of soldier production might vary between different clones.

**Significance Statement:** Nest-living has often been considered to be a necessary condition for the evolution of eusociality. In a small number of aphid species, however, sterile soldier castes have evolved in open colonies without a nest. To understand why these aphids are unique, we examined the kin structure and genetic relatedness of individuals within eusocial and non-eusocial open colonies of two aphid species on bamboo. We found that clonal mixing occurred in both species, but the eusocial colonies are more genetically homogenized, functioning as a large colony. Our results suggest that ecological conditions that promote genetically homogenized, large and long-lived colonies are important for the evolution of eusociality in these aphids. We propose that the open colonies of social aphids provide an ideal model system in which to study the evolution of altruism.

## 1. Introduction

Understanding how complex sociality has evolved and is maintained is a fundamental question in evolutionary biology. Two key factors need to be addressed: the costs and benefits of helping and being helped, and the genetic relatedness of the actors. The mode of reproduction, such as monogamy or parthenogenesis, can enhance genetic relatedness (Hughes et al. 2008; Boomsma 2009), and the ecological context, such as the intensity of parasitism or the availability of food, can influence the benefit-to-cost ratio of helping behaviour (Korb and Heinze 2008; Korb and Heinze 2016). Living in a nest can influence both of these factors (Starr, 1991). Confinement in a structure may make colony defence a more viable proposition; the structure itself, for example a plant gall, might provide a food source; and the construction and maintenance of the structure might select for adaptations for altruism in the colony members (Stone and Schönrogge 2003). Independently, enclosure in a structure will tend to keep close kin together and potentially increase relatedness within the colony. Therefore, although the role of the nest in the lives of social insects is complex and nuanced (McGlynn, 2012; Bengston and McGlynn, 2021), most authors consider that nest-living is a necessary condition for the initial evolution of eusociality, in both fortress defenders (for example aphids) and life-insurers (such as ants) (e.g. Foster and Northcott 1994; Queller and Strassmann 1998; Hölldobler and Wilson 2009).

Within the aphids, social behaviour has evolved only within those clades that were ancestrally gall-formers on plants (Foster and Northcott 1994; Stern 1994; Stern 1998). The gall is induced by the feeding behaviour of the aphid foundress, whose clonal descendants are provided with a defendable fortress that is also a source of food. Gall-living provides a selective context for the evolution of altruistic behaviour such as gall-cleaning (removing honeydew) (Aoki 1980, Benton and Foster 1991, Kutsukake et al. 2012, Uematsu et al. 2018, Kutsukake et al. 2019a, Shibao et al 2021), gall repair (Kurosu et al. 2003, Pike and Foster 2004, Kutsukake et al. 2009, Kutsukake et al. 2019b) and, in some but not all species, colony defence. In about 40 out of the 5,000 species of aphids, a morphologically specialized and sterile soldier caste has evolved (Pike and Foster 2008; Aoki and Kurosu 2010; Abbot and Chapman 2017). These species are eusocial, showing overlap of generations, cooperative care of the young, and reproductive altruism (Costa 2006).

Arguably the most puzzling of all the eusocial aphids are those species that have sterile soldiers and live in open colonies on plant surfaces without modifying their structures. These are the only known eusocial animals that do not occupy some kind of nest. Sterile soldiers in open colonies have evolved at least twice independently, in the subfamilies Eriosomatinae and Hormaphidinae (Stern and Foster 1996; Abbot 2015). In both clades, the origin of eusocial behaviour in open colonies on the secondary host is considered to be evolutionarily independent from the origin of eusociality in the aphid generations that live in galls on the primary tree host, where sexual reproduction occurs (Stern 1998). Very little is known about either the genetic relatedness between colony members or the costs and benefits of helping behaviour within these open aphid colonies. Most of the few studies of clonal mixing in relation to sociality in aphids have focussed on the gall generations. These studies have shown that clonal mixing does occur within the galls, that the level of mixing is very variable, and that the dispersing aphids can adjust their social and reproductive behaviour depending on whether they are in their own gall or that of another clone (Abbot et al. 2001; Johnson et al. 2002; Wang et al. 2008; Abbot 2009; Miller et al. 2015). By contrast, there has been only one study of the genetic structure of soldier-producing aphids in open colonies (Hattori et al. 2015). This showed that there was some clonal mixing in five open colonies of *Ceratovacuna japonica* using AFLP markers. But this study was on a very small scale (about 10 m x 5 m), much smaller than the dispersal distance of individual aphids, and of a small sample size. Further genetic analysis of the open colonies of soldier-producing aphids on a wider spatial scale is needed to reveal the clonal diversity in a population, the kin structure and genetic relatedness within a colony, and the evolutionary forces that determine the proportion of sterile soldiers.

The aim of our study is to use newly developed microsatellite markers to provide a detailed account of the genetic relatedness amongst individual social aphids living in open colonies. We studied two cerataphidine species with contrasting levels of sociality on the secondary host: *Astegopteryx bambucifoliae* (Takahashi), which is non-eusocial (Figure 1a, c); and *Pseudoregma alexanderi* (Takahashi), which is eusocial (Figure 1b and d). Both species belong to genera that have eusocial colonies on the primary woody host (*Styrax*) and that form colonies on the secondary host (bamboos etc). The crucial difference between the two species is that aphids of the genus *Pseudoregma* (including *P. alexanderi*) living in open colonies on the secondary host have evolved a novel kind of sterile first-instar horned soldier that defends the colonies, which can therefore be described as eusocial on the secondary host: aphids of the genus *Astegopteryx* (including *A. bambucifoliae*) have not evolved specialized sterile soldiers on the secondary host (Figure S1). Both species were studied on the same secondary host, the giant bamboo *Dendrocalamus latiflorus*.

**Figure 1.**
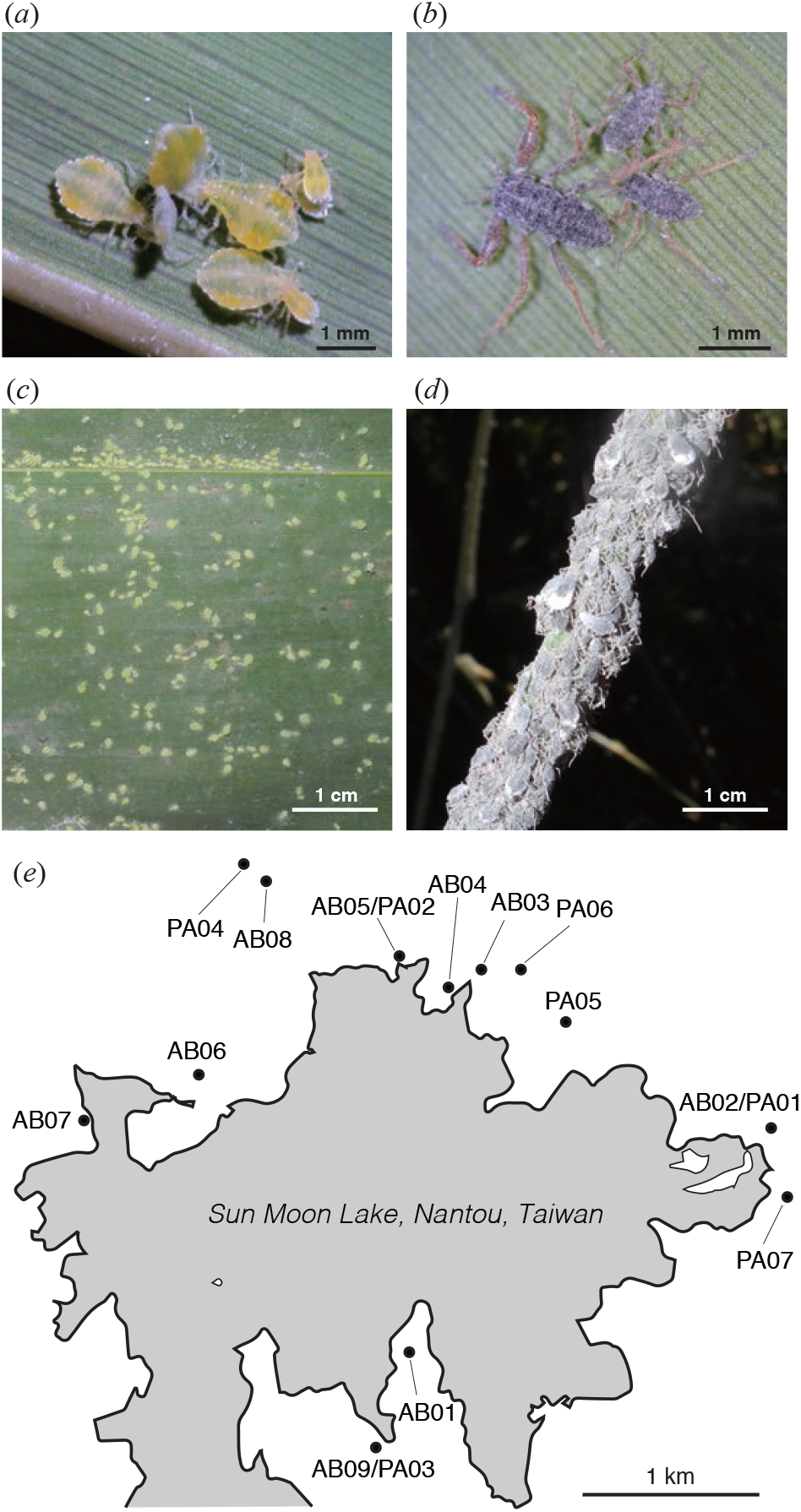
Study species and sampling sites. (*a*) *Astegopteryx bambucifoliae*. (*b*) *Pseudoregma alexanderi*. A soldier nymph (left) and normal first-instar nymphs (right). (*c, d*) A colony of *A. bambucifoliae* (*c*) and *P. alexanderi* (*d*). (*e*) Sampling sites of *A. bambucifoliae* (AB01-AB09) and *P. alexanderi* (PA01-PA07). A dot shows the location of a bamboo clump where aphids were collected.

The key questions we intend to answer here are: (i) what is the level of clonal mixing in these open colonies of social aphids; (ii) does the level of clonal mixing correlate with the level of aphid sociality; (iii) what can we infer from our observations about how sociality may have evolved and be maintained in the absence of nest-living?

## 2. Materials and methods

### 2.1. Study species

#### Astegopteryx bambucifoliae

This aphid forms eusocial colonies in galls, which include sterile second-instar soldiers and may contain up to 5,000 aphids, on the primary woody host *Styrax suberifolia* (Kurosu and Aoki 1991). Viviparous winged aphids disperse from the galls from August to November to form open colonies on the undersides of the leaves of their secondary host plants, *Bambusa* spp. (*dolichoclada, edulis, oldhami, stenostachya*), *Dendrocalamus latiflorus*, and *Phyllostachys lithophia* (Liao 1976).

The open colonies of this aphid on the secondary host are not eusocial. None of the early-instar nymphs are sterile and we never observed them attacking predators, although all instars can use their frontal horns to attack each other for feeding sites (Aoki and Kurosu 1985). There is one brief report of soldier behaviour in *Astegopteryx bambucifoliae*: a total of eight first-instar aphids were observed to have the tips of their rostra in contact either with some unidentified insect eggs or scymnine (Coccinellidae) pupae, but no evidence was presented that the stylets had penetrated (Aoki and Kurosu 1989). These observations are suggestive, but do not provide robust support that this species has altruistic defensive behaviour that deters or kills predators.

Winged individuals emerge from bamboo colonies from February to May and disperse to *Styrax suberifolia* to give rise to the females and males that produce eggs, but a small number of aphids remain on bamboo and continue parthenogenetic reproduction (Takahashi 1923).

#### Pseudoregma alexanderi

Most species of this genus form eusocial colonies on the primary host (*Styrax* spp.) with sterile second-instar soldiers, as in *Astegopteryx* (Aoki and Kurosu 2010). This species has been found in Taiwan (Liao 1976), China (Zhang et al. 2013), Philippines (Calilung, 1985), Nepal (Sharma 1968) and India (Ghosh 1988) on bamboos. However, galls of *P. alexanderi* have never been recorded on a primary host plant. In contrast to *A. bambucifoliae*, therefore, *P. alexanderi* in Taiwan is permanently confined to the secondary host plant, *Dendrocalamus latiflorus* (Figure S1).

The open colonies of *P. alexanderi* on the shoots and stems of the secondary host, where wingless viviparous females reproduce throughout the year, are eusocial. The clones produce two types of first-instar nymphs: sterile soldiers and normal fertile nymphs. The soldiers are distinguishable from the normal nymphs of the same instar by their larger body size, sclerotized exoskeletons, enlarged and thickened forelegs, and well-developed sharp frontal horns, and they do not moult and grow to adulthood (Aoki 1978). They defend their colonies by clasping predators with enlarged forelegs and piercing their skins with a pair of sharp horns on a head (Aoki et al. 1981). The horns and piercing behaviour of soldiers might be derived from head-butting behaviour between colony members, which is found in almost all of the cerataphidine aphids on the secondary host (Aoki and Kurosu 1985; Foster 1996; Howard et al. 1998; Morris and Foster 2008). The soliders in a colony of *P. alexanderi* shows bimodal distribution for morphological characters (Aoki and Miyazaki 1978, Stern et al. 1996). First-instar normal nymphs can disperse by being carried on the wind (Aoki et al. 1981). In spring, winged adults that give rise to the females and males (sexuparae) emerge but they have never been recorded on a primary host.

### 2.2. Sampling

Both aphid species were sampled at Sun Moon Lake, Nantou, Taiwan: *A. bambucifoliae* on 18 November 2012 and *P. alexanderi* on 18 November 2012 and 23 April 2014. The host plant *D. latiflorus* forms a dense cluster of stems (a clump). Aphids were collected from nine (AB01-AB09) and seven (PA01-04 in 2012 and PA05-07 in 2014) clumps of *D. latiflorus* for *A. bambucifoliae* and *P. alexanderi* respectively over an area of more than 1,000 hectares (Figure 1e). In the present study, we defined a “colony” as a group of aphids on the same leaf for *A. bambucifoliae* and on the same developing shoot for *P. alexanderi*, in order to facilitate comparison between the two species. For each clump, two colonies were collected from different stems, except for clump PA04, in which only one colony of *P. alexanderi* was collected. In total, we collected 18 colonies of *A. bambucifoliae* and 13 of *P. alexanderi*. The numbers of aphids in the colonies that we sampled were roughly similar in the two species, ranging from 52 to 882 (mean 344.5) in *A. bambucifoliae* and from 56 to 831 (mean 235.3) in *P. alexanderi*. Note that it was not possible to record data blind because our study involved focal animals in the field.

All the aphids from each colony were put into 95% ethanol and the total number of aphids in each colony was counted under a dissecting microscope. For *P. alexanderi*, the numbers of soldier first-instar nymphs were also counted for each colony.

From each colony, 15 – 24 individuals from four age classes (first-instar, second or third instar, third or fourth instar, and wingless adult in *A. bambucifoliae*; first-instar soldier, first-instar normal, from second to fourth instar, and wingless adult in *P. alexanderi*) were selected for DNA extraction and genotyping. In total, 364 individuals of *A. bambucifoliae* and 251 of *P. alexanderi* were genotyped. In addition, from two clumps (PA03 and PA07), where we found multiple clones of *P. alexanderi* by the first genotyping, 112 first-instar (57 soldier and 55 normal) nymphs were subsequently genotyped without breaking their exoskeleton to associate morphological characters with a genotype.

### 2.3. DNA extraction

DNA was extracted using a glass milk extraction method in a microtitre plate as described in Amos et al. (2016). In the first genotyping, each aphid was crushed with a plastic pestle or a pipette tip in 40 μl of lysis solution (10 mM Tris HCl (pH 8.0), 1 mM EDTA, 1% SDS) with proteinase K and incubated at 56°C for 12 hours. The DNA was adsorbed to flint glass particles in the presence of a 3× excess of 6 M NaI. After two ethanol washes, the DNA was eluted in 50 µl of low TE buffer. In the additional genotyping in which the exoskeleton was not crushed, we punctured the body of a nymph with an insect pin and incubated it in the lysis solution with proteinase K described above at 56°C for 12 hours to extract DNA and clear inside the body. After three hours, the exoskeleton was picked up and the remaining solution was subjected to DNA extraction and genotyping. The exoskeleton was examined as described below in section 2.6.

### 2.4. Microsatellite isolation

Extracted DNA from a wingless adult of *A. bambucifoliae* and *P. alexanderi* were sequenced using Illumina Miseq. From the sequence data, we searched for microsatellites using custom C++ scripts, focusing on two motifs: (AC)_n_ and (AT)_n_ as described in (Amos and Filipe 2014). Then we used the web software Websat (Martins et al. 2009) to design primer sequences for each candidate microsatellite. We tested each locus for polymorphism using panels of 8-16 individuals from different colonies, and successfully isolated eight primer pairs for *A. bambucifoliae* and nine primer pairs for *P. alexanderi* that showed polymorphism.

### 2.5. Genotyping

Polymerase Chain Reactions (PCRs) were performed using Type-it Microsatellite PCR Kit (Qiagen) in 10 µl reactions. PCR conditions were as follows: an initial denaturing step of 5 min at 95°C, followed by 35 cycles of 95°C denaturation for 30 sec, 60°C annealing for 90 sec, 72°C extension for 60 sec, followed by a final extension step for 30 min at 60°C. One primer from each pair was labelled with a fluorescent dye (6-FAM / HEX / NED) as described in Table 1. After amplification, the PCR product was mixed with The GeneScan 500 LIZ Size Standard (Applied Biosystems) and subjected to fragment analysis. Allele calling was performed by GeneMapper software (Applied Biosystems) and checked manually. Samples with ambiguous genotype were re-run twice and the consensus genotype was taken forward for analysis. To reduce the possible impact of genotyping errors / *de novo* mutations, two individuals with unique multilocus genotypes that were found only once and differed from another genotype by only one single allele were excluded from the analysis. In total, 360 individuals of *A. bambucifoliae* and 244 of *P. alexanderi* were subjected to further statistical analysis.

**Table 1.**
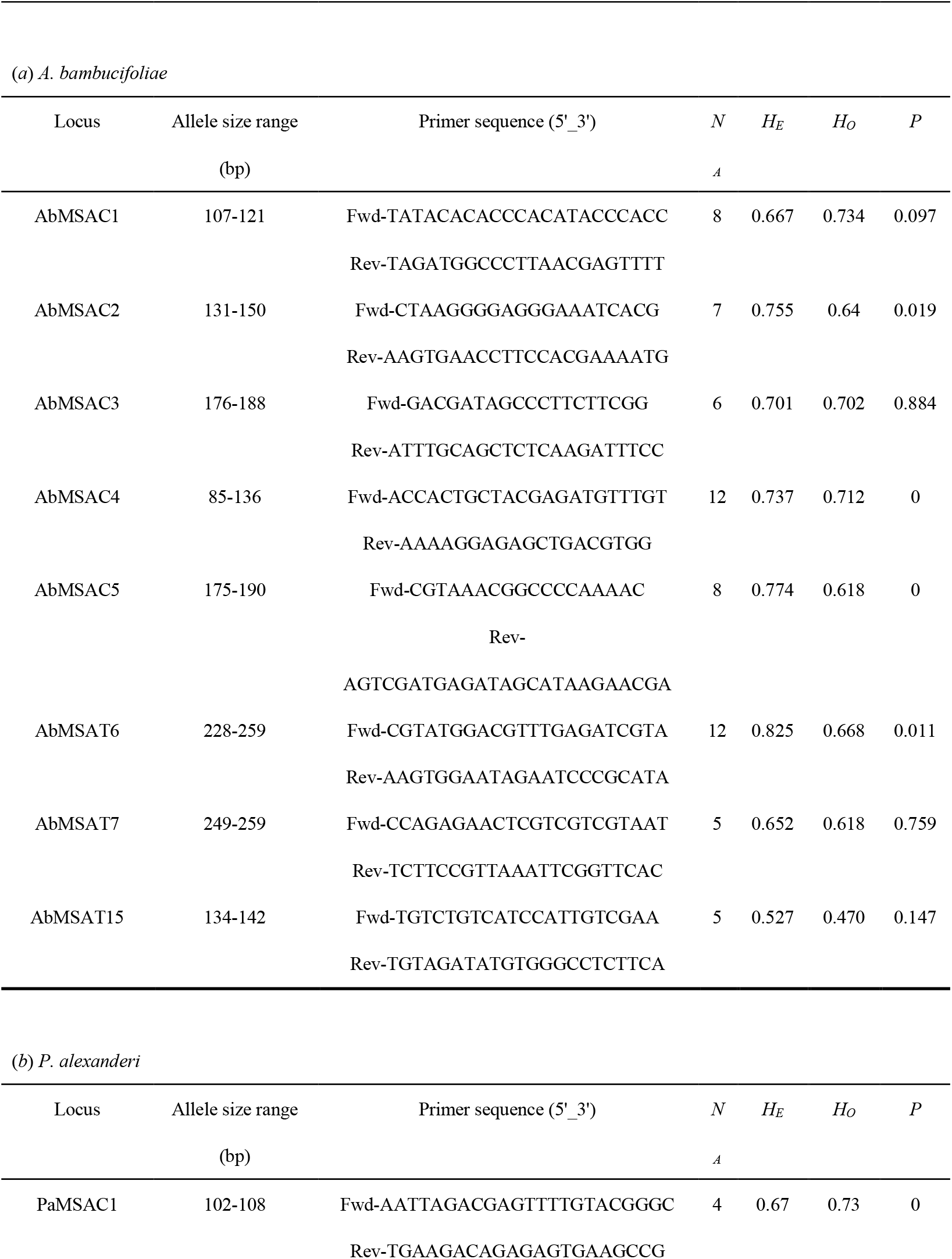
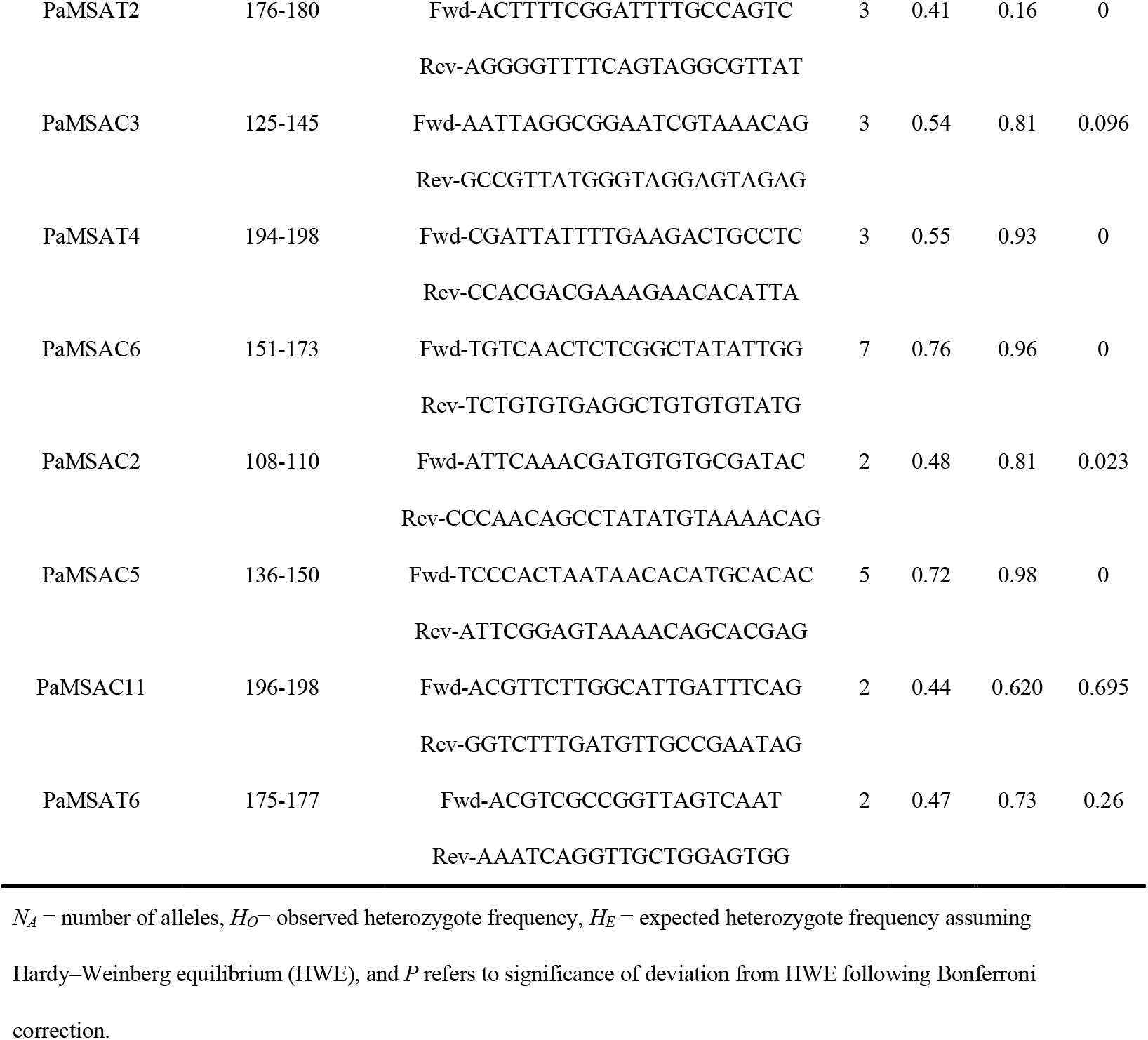
Summary statistics of the polymorphic microsatellite loci with the primer sequences in (*a*) *A. bambucifoliae* and (*b*) *P. alexanderi*.

### 2.6. Morphological examination of first-instar nymphs in

***P. alexanderi*** Exoskeletons picked up from the proteinase K solution were washed with distilled water and mounted on a slide using Mount-quick Aqueous (Daido Sangyo, Japan). The mounted individuals were photographed using a digital camera (Nikon 1 V2) attached to a light microscope. Their morphological characters (body length, fore-femur length, horn length) were measured using ImageJ software (Schneider et al., 2012).

### 2.7. Data analysis

All statistical analyses were conducted using the software R (R Core Team 2022). Different clones were identified using their multilocus genotypes (MLGs) and the R package *poppr* (Kamvar et al. 2013). Deviations from Hardy-Weinberg Equilibrium (HWE) at each locus were tested based on 1000 Monte Carlo permutations of alleles implemented in R package *pegas* v0.9 (Paradis 2010), using only one individual per MLG. Following others (Gilbert et al. 2007; Vantaux et al. 2011; Kronauer et al. 2013), we use mean pairwise relatedness as a measure both of the potential for altruism and of the genetic similarity between groups of aphids. Pairwise relatedness values among all individuals were calculated following Queller and Goodnight (Queller and Goodnight 1989) using Kingroup (Konovalov et al. 2004) software. Since the pairwise data are non-independent, we could not directly compare average relatedness values between the different age or morph categories (Prugnolle and de Meeus 2002). Instead, we performed permutation tests (Coulon et al. 2006) by generating 1000 random resampling sets without replacement, such that each individual occurs only once in a given resampled set. Resampled individuals were chosen from the same colony. Under the null hypothesis, the difference of mean relatedness should follow a normal distribution centred on zero; under the alternative hypothesis (i.e. category-biased dispersal), the difference should be significantly different from zero. SPAGeDI (Hardy and Vekemans 2002) software was used to test whether the pairwise regression relatedness among the aphids decreased with the distance between the aphids. An analysis of molecular variance (AMOVA) (Excoffier et al. 1992) was performed to test for population differentiation in order to show if a pair of colonies on a bamboo clump are demographically distinct. If significant variation occurs between the two colonies, then we inferred that they are genetically distinct units. For the morphological characters of soldier nymphs in *P. alexanderi*, bimodality was tested using Hartigan’s dip test (Hartigan and Hartigan 1985) implemented in the R package *diptest*.

## 3. Results

### 3.1. Size and disposition of the aphid colonies

Both of the aphid species feed on the same host plant, the Taiwan giant bamboo *Dendrocalamus latiflorus*, but the size and disposition of the colonies of the two species are markedly different. *A. bambucifoliae* feeds on the underside of the bamboo leaves, forming relatively small, discrete colonies (Figure 1c), whereas *P. alexanderi* feeds on shoots and stems, where the individual colonies tend to coalesce into a large colony, often of immense size (Figure 1d). Although we did not measure aphid density, measurements on a related species with similar habits (*Pseudoregma baenzigeri*) suggest there would be around 10 adults or 120 first-instar nymphs per square cm, equivalent to 28,000 adults or 340,000 first-instar nymphs for every metre of 9-cm diameter bamboo stem (Aoki et al. 2007). The colonies that we studied were at this, or even greater, densities.

### 3.2. Clonal diversity and genetic differentiation

The number of alleles and the heterozygosities (observed and expected) for each locus are given in Table 1. The number of alleles per locus ranged from 5 to 12 (mean 7.9) in *A. bambucifoliae* and from 2 to 7 (mean 3.4) in *P. alexanderi*. When using only one individual per MLG, deviations from Hardy-Weinberg equilibrium (HWE) were significant (*P* < 0.05) in four of eight loci in *A. bambucifoliae* and in six of nine loci in *P. alexanderi*. A total of 99 different clones were recognized based on MLGs by using eight microsatellite markers in the study population of *A. bambucifoliae* and 19 different clones by using nine markers in *P. alexanderi*. In *A. bambucifoliae*, each clone was restricted to a single sampling site. By contrast, in *P. alexanderi* three clones were found at multiple (two to six) widely dispersed sites. 94% (230/244) of the genotyped individuals of *P. alexanderi* consisted of the four major clones, described as “A”, “B”, “C”, and “D” (Table 2). 89% (16/18) of the *A. bambucifoliae* colonies and 77% (10/13) of the *P. alexanderi* colonies consisted of multiple clones (two to 20 clones in *A. bambucifoliae*, two to six in *P. alexanderi*), although half (5/10) of the mixed-clone *P. alexanderi* colonies and 25% (4/16) of the mixed-clone *A. bambucifoliae* colonies were almost monoclonal, with more than 80% of the colony members being of a single genotype.

**Table 2.**
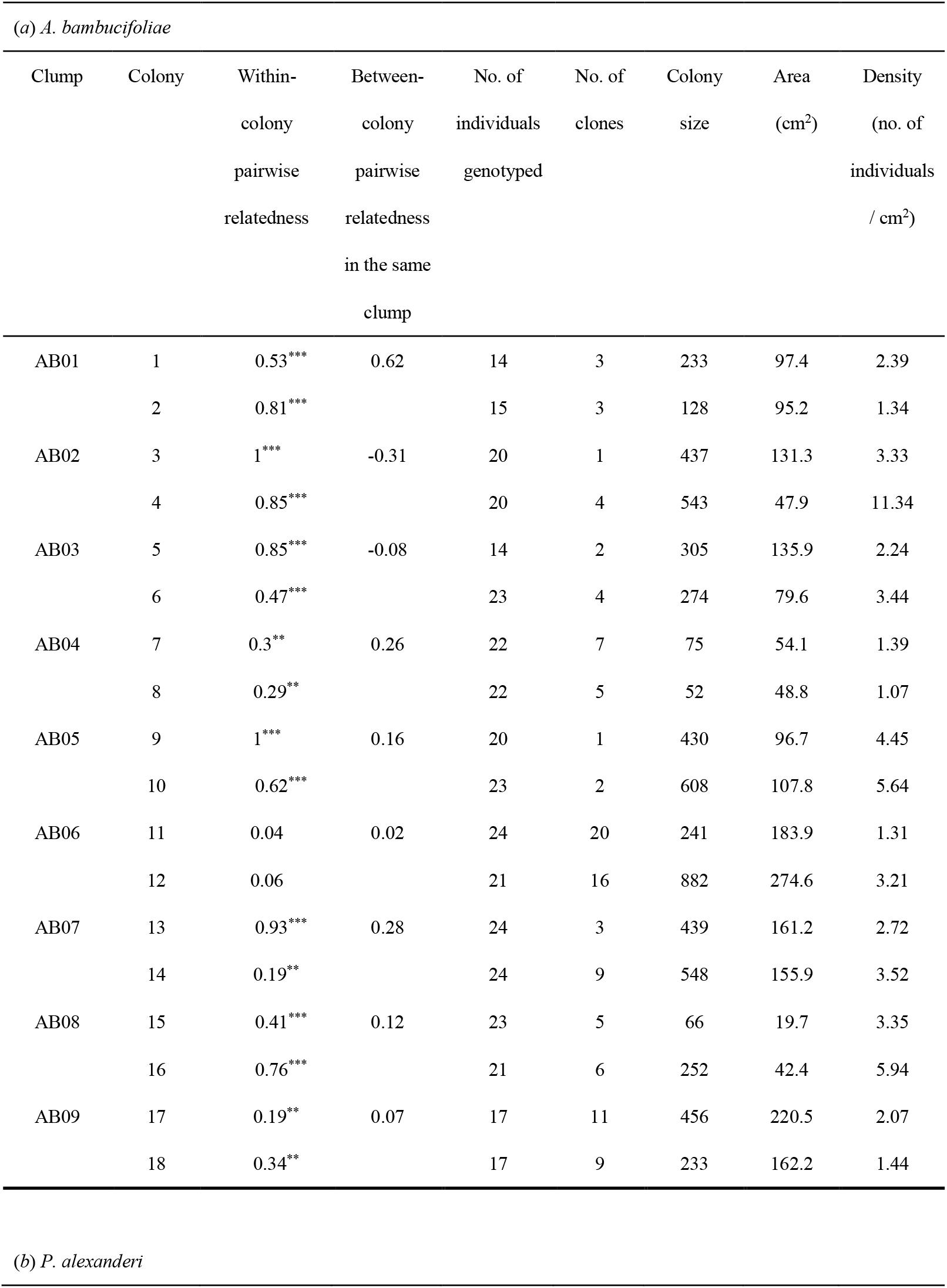
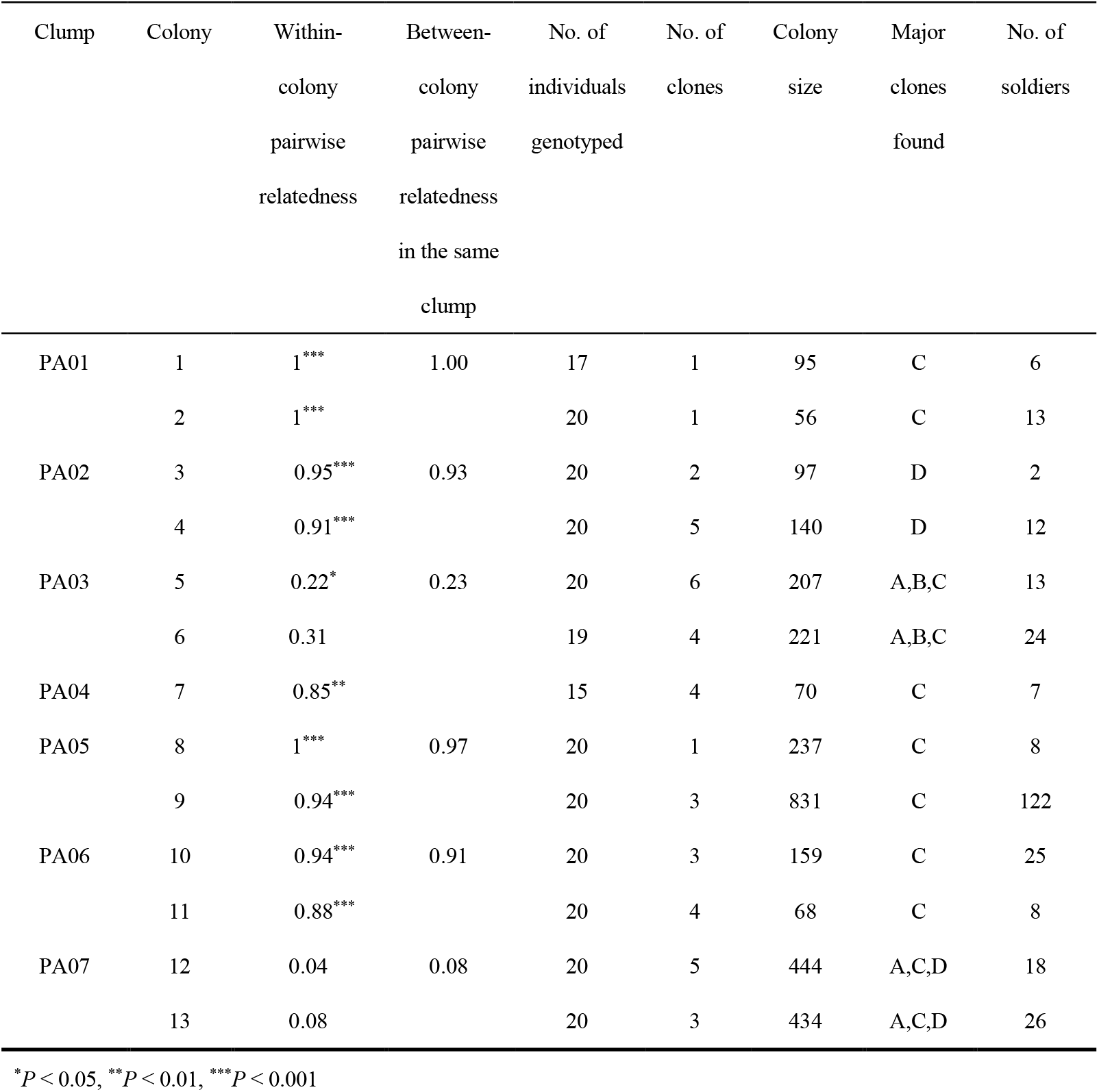
Colony structure of (*a*) *A. bambucifolia*e and (*b*) *P. alexanderi* colonies that were collected and partly genotyped.

| (a) *A. bambucifoliae* |  |  |  |  |  |  |  |  |
| --- | --- | --- | --- | --- | --- | --- | --- | --- |
| Clump | Colony | Within-colony pairwise relatedness | Between-colony pairwise relatedness in the same clump | No. of individuals genotyped | No. of clones | Colony size | Area (cm<sup>2</sup>) | Density (no. of individuals / cm<sup>2</sup>) |
| AB01 | 1 | 0.53*** | 0.62 | 14 | 3 | 233 | 97.4 | 2.39 |
|  | 2 | 0.81*** |  | 15 | 3 | 128 | 95.2 | 1.34 |
| AB02 | 3 | 1*** | -0.31 | 20 | 1 | 437 | 131.3 | 3.33 |
|  | 4 | 0.85*** |  | 20 | 4 | 543 | 47.9 | 11.34 |
| AB03 | 5 | 0.85*** | -0.08 | 14 | 2 | 305 | 135.9 | 2.24 |
|  | 6 | 0.47*** |  | 23 | 4 | 274 | 79.6 | 3.44 |
| AB04 | 7 | 0.3** | 0.26 | 22 | 7 | 75 | 54.1 | 1.39 |
|  | 8 | 0.29** |  | 22 | 5 | 52 | 48.8 | 1.07 |
| AB05 | 9 | 1*** | 0.16 | 20 | 1 | 430 | 96.7 | 4.45 |
|  | 10 | 0.62*** |  | 23 | 2 | 608 | 107.8 | 5.64 |
| AB06 | 11 | 0.04 | 0.02 | 24 | 20 | 241 | 183.9 | 1.31 |
|  | 12 | 0.06 |  | 21 | 16 | 882 | 274.6 | 3.21 |
| AB07 | 13 | 0.93*** | 0.28 | 24 | 3 | 439 | 161.2 | 2.72 |
|  | 14 | 0.19** |  | 24 | 9 | 548 | 155.9 | 3.52 |
| AB08 | 15 | 0.41*** | 0.12 | 23 | 5 | 66 | 19.7 | 3.35 |
|  | 16 | 0.76*** |  | 21 | 6 | 252 | 42.4 | 5.94 |
| AB09 | 17 | 0.19** | 0.07 | 17 | 11 | 456 | 220.5 | 2.07 |
|  | 18 | 0.34** |  | 17 | 9 | 233 | 162.2 | 1.44 |
| (b) *P. alexanderi* |  |  |  |  |  |  |  |  |

| Clump | Colony | Within-colony<br>pairwise<br>relatedness | Between-colony<br>pairwise<br>relatedness<br>in the same<br>clump | No. of<br>individuals<br>genotyped | No. of<br>clones | Colony<br>size | Major<br>clones<br>found | No. of<br>soldiers |
| --- | --- | --- | --- | --- | --- | --- | --- | --- |
| PA01 | 1 | 1<sup>***</sup> | 1.00 | 17 | 1 | 95 | C | 6 |
|  | 2 | 1<sup>***</sup> |  | 20 | 1 | 56 | C | 13 |
| PA02 | 3 | 0.95<sup>***</sup> | 0.93 | 20 | 2 | 97 | D | 2 |
|  | 4 | 0.91<sup>***</sup> |  | 20 | 5 | 140 | D | 12 |
| PA03 | 5 | 0.22<sup>*</sup> | 0.23 | 20 | 6 | 207 | A,B,C | 13 |
|  | 6 | 0.31 |  | 19 | 4 | 221 | A,B,C | 24 |
| PA04 | 7 | 0.85<sup>**</sup> |  | 15 | 4 | 70 | C | 7 |
| PA05 | 8 | 1<sup>***</sup> | 0.97 | 20 | 1 | 237 | C | 8 |
|  | 9 | 0.94<sup>***</sup> |  | 20 | 3 | 831 | C | 122 |
| PA06 | 10 | 0.94<sup>***</sup> | 0.91 | 20 | 3 | 159 | C | 25 |
|  | 11 | 0.88<sup>***</sup> |  | 20 | 4 | 68 | C | 8 |
| PA07 | 12 | 0.04 | 0.08 | 20 | 5 | 444 | A,C,D | 18 |
|  | 13 | 0.08 |  | 20 | 3 | 434 | A,C,D | 26 |

\* $P < 0.05$ , \*\* $P < 0.01$ , \*\*\* $P < 0.001$

### 3.3 Relatedness within and between the colonies on a single clump

Average pairwise relatedness between individuals within a colony ranged from 0.04 to 1.0 (mean = 0.54, n = 18) in *A. bambucifoliae* and from 0.04 to 1.0 (mean = 0.71, n = 13) in *P. alexanderi*, which is not significantly different between the two species (Welch’s t-test, *t* = -1.27, df = 23.4, *P* = 0.22, Figure 2 and Table 2). The relatedness value is significantly higher than predicted by a random distribution in 89% (16/18) of *A. bambucifoliae* colonies and 77% (10/13) colonies of *P. alexanderi* (permutation test, *P* < 0.05, Table 2). Average pairwise relatedness between individuals in the different colonies of the same clump ranged from -0.31 − 0.62 (mean = 0.13, n = 9) in *A. bambucifoliae* and from 0.07 − 1.0 (mean = 0.69, n = 6) in *P. alexanderi*, which is significantly different between the two species (Welch’s t-test, *t* = -2.94, df = 7.6, P = 0.02, Figure 3 and Table 2).

**Figure 2.**
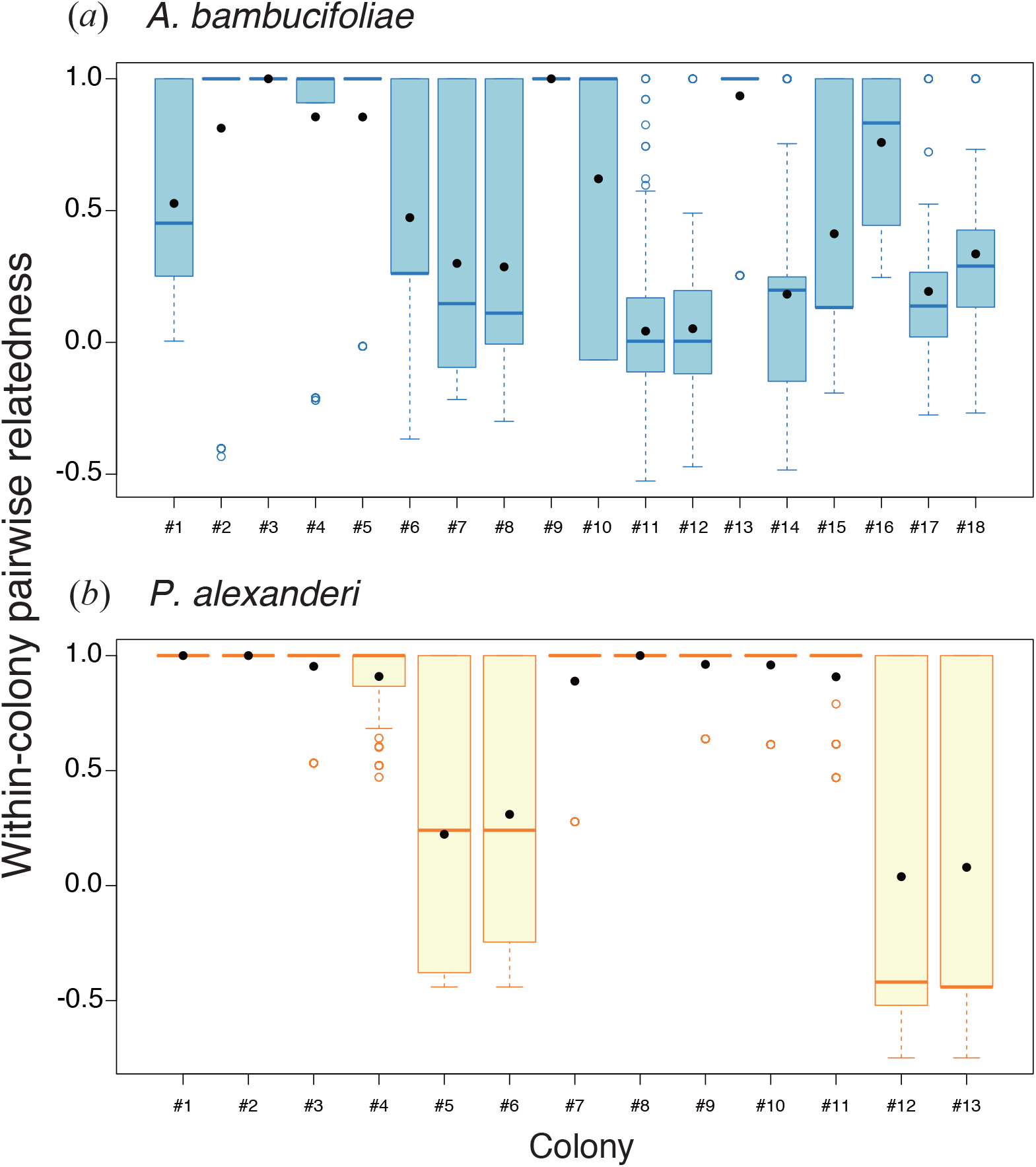
Within-colony pairwise genetic relatedness (n = 18 for *A. bambucifoliae* (*a*); n = 13 for *P. alexanderi* (*b*)). Box plots show the range of values, the first and third quartiles and medians. Open circles show outliers and filled circles show means.

**Figure 3.**
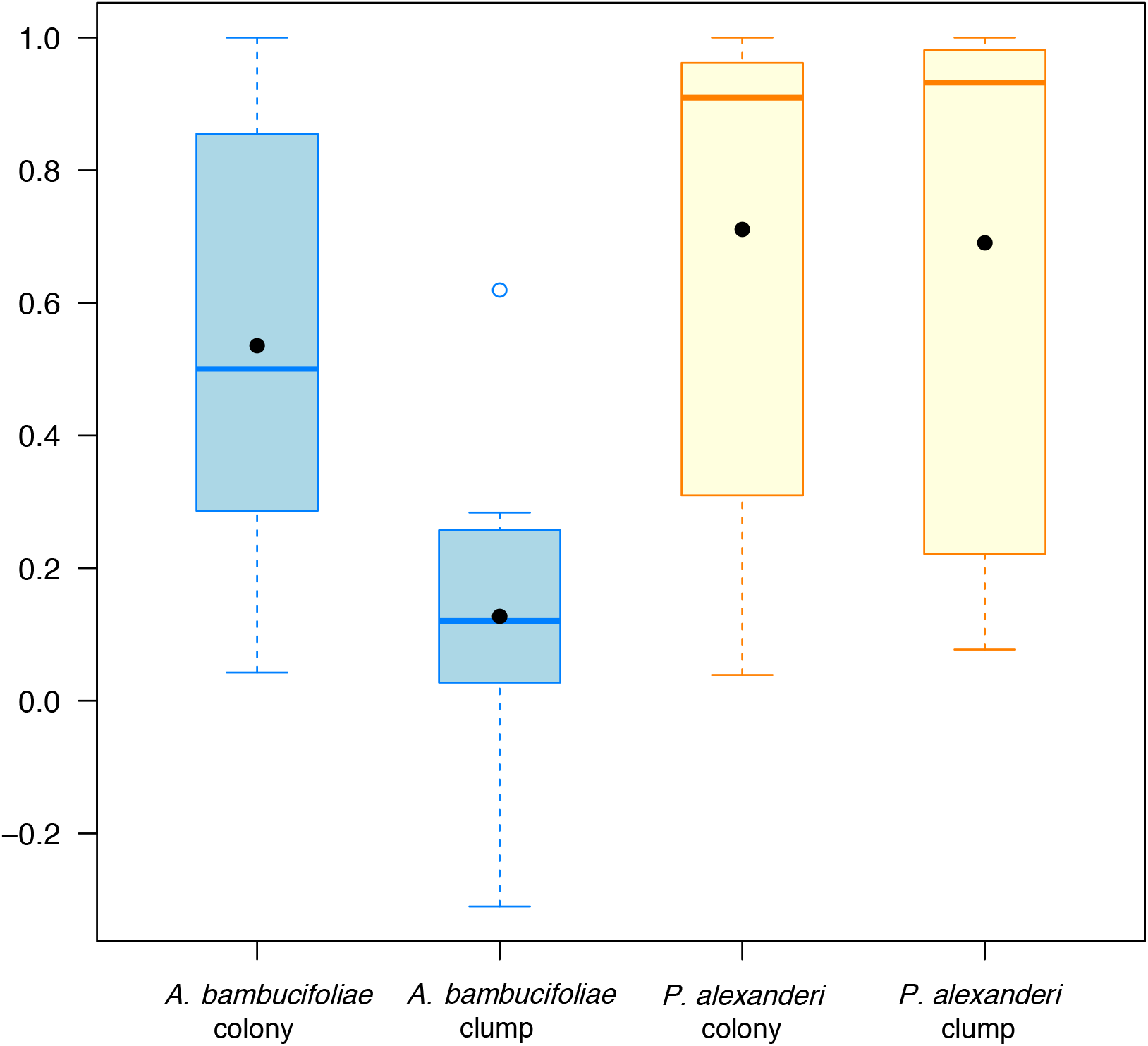
Average values of within-colony relatedness and between-colony relatedness in the same clump in *A. bambucifoliae* and *P. alexanderi*. Box plots show the range of values, the first and third quartiles and medians. Open circles show outliers and filled circles show means.

Table 3 reveals significant variation in genetic variance among different bamboo clumps (16%) and among different colonies on the same clump (38%) in *A. bambucifoliae*, indicating a dispersal process in which related colonies do not usually end up as neighbours. By contrast, in *P. alexanderi*, there was significant variation in genetic variance among different bamboo clumps (63%) but not among different colonies on the same clump (1.4%), suggesting much more limited dispersal and a tendency for colonies to grow and split within a clump. If dispersal is primarily associated with one age class and if dispersers occasionally enter an unrelated colony, we would expect that age class to exhibit reduced pairwise relatedness. In fact, average pairwise relatedness was not significantly different among pairs of the four age-classes in *A. bambucifoliae* (permutation test, *P* > 0.05; electronic supplementary material, Figure S2). The SPAGeDI analysis showed that pairwise relatedness did not decrease significantly with distance between samples for either species (*A. bambucifoliae*: *r*^2^ = 0.0007, y = -0.03 - 0.007x, one-tailed *P* = 0.34; n = 360 individuals and 57470 pairs; *P. alexanderi*: *r*^2^ = 0.0002, y = -0.15 + 0.01x, one-tailed *P* = 0.48; n = 245 individuals and 25439 pairs), suggesting that dispersal distances are either much smaller or much larger that the spatial scale of sampling in our study.

**Table 3.** Analysis of molecular variance (AMOVA) table for populations of (*a*) *A. bambucifoliae* and (*b*) *P. alexanderi*.

| Sources of variation | df | % of variation | *Phi*-statistics | *P*<sup>a</sup> |
| --- | --- | --- | --- | --- |
| Between tree | 8 | 15.8 | 0.158 | 0.003 |
| Between colonies within tree | 9 | 37.7 | 0.447 | < 0.001 |
| Within colonies within tree | 342 | 46.5 | 0.535 |  |

(b) *P. alexanderi*
| Sources of variation | df | % of variation | Phi-statistics | *P*<sup>a</sup> |
| --- | --- | --- | --- | --- |
| Between tree | 6 | 62.8 | 0.628 | 0.001 |
| Between colonies within tree | 6 | 1.4 | 0.038 | 0.123 |
| Within colonies within tree | 232 | 35.8 | 0.642 |  |

<sup>a</sup>*P*-values are based on 9999 permutations.

### 3.4 Differences in the proportion and morphology of soldiers among genotypes in *P. alexanderi*

To test whether different clones were more or less likely to produce soldiers, we additionally genotyped 57 soldier and 55 normal first instar nymphs from the two clumps, PA03 and PA07. When included with 25 soldiers and 20 normal first instar nymphs that had already been genotyped, 96% (150/157) of the genotyped nymphs consisted of four major clones, “A”, “B”, “C”, and “D” described above, and all of the four clones contained soldiers. However, the proportion of soldiers to normal first-instar nymphs was significantly different among the four clones in both of the clumps (Fisher’s exact test, *P* = 0.006 in PA03 and *P* = 0.01 in PA07, Table 4). For the two clones (“A” and “C”) that occurred in both of the clumps, fewer soldiers of genotype A and more of genotype C were found than was expected (Table 4).

**Table 4.**
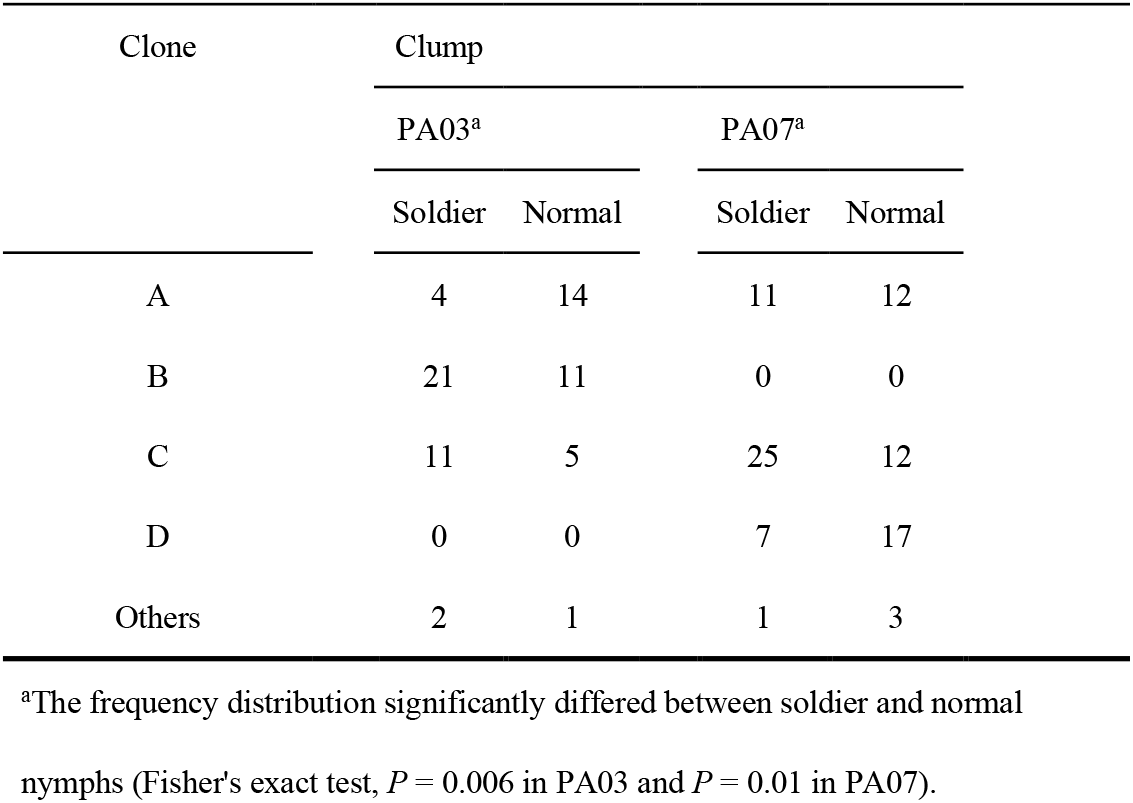
Clonal composition of first-instar nymphs in the two clone-mixed clumps.

| Clone | Clump |  |  |  |
| --- | --- | --- | --- | --- |
|  | PA03<sup>a</sup> |  | PA07<sup>a</sup> |  |
|  | Soldier | Normal | Soldier | Normal |
| A | 4 | 14 | 11 | 12 |
| B | 21 | 11 | 0 | 0 |
| C | 11 | 5 | 25 | 12 |
| D | 0 | 0 | 7 | 17 |
| Others | 2 | 1 | 1 | 3 |

<sup>a</sup>The frequency distribution significantly differed between soldier and normal nymphs (Fisher's exact test, $P = 0.006$ in PA03 and $P = 0.01$ in PA07).

To test whether the dimorphism of soldiers of *P. alexanderi*, shown in previous studies (Aoki and Miyazaki 1978, Stern et al. 1996), is genetically or environmentally determined, we performed Hartigan’s dip test (Hartigan and Hartigan 1985) implemented in the R package *diptest* using the morphological characters of the genotyped soldier nymphs. The fore-femur length of the soldiers in both of the two mixed-clone clumps was bimodal, as judged by Hartigan’s dip test: (*P* = 0.006 in clump PA03 and *P* = 0.003 in clump PA07, Figure 4). Horn length was also bimodal, but the dip test is only significant in clump PA07 (*P* = 0.42 in PA03 and *P* = 0.01 in PA07, Figure 4). Even when analysed separately for each clone, fore-femur and horn length were bimodal in most clones, except for “A” in PA03 and “D” in PA07 (Figure S3).

**Figure 4.**
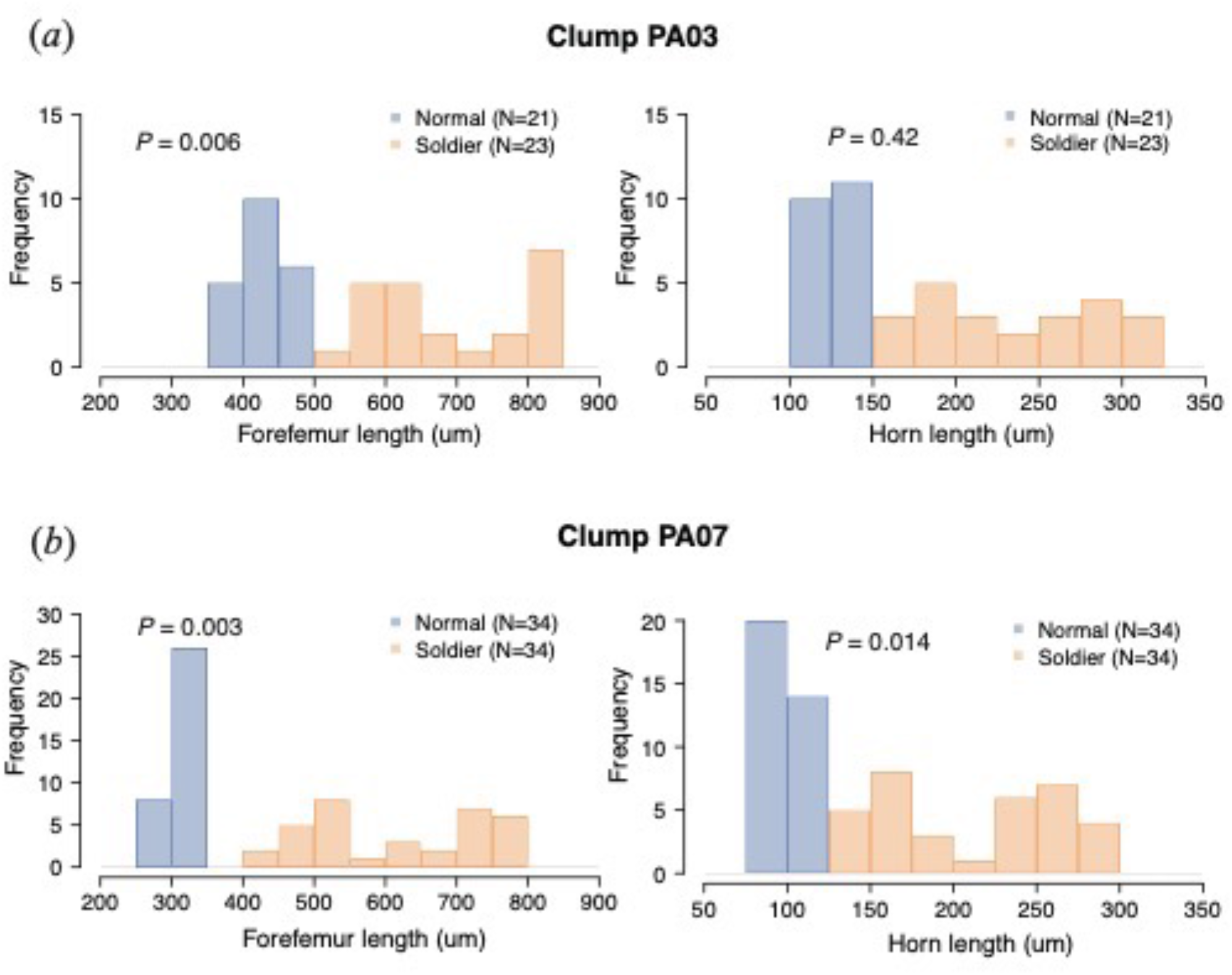
Frequency distributions of the morphological characters (horn length and fore-femur length) in the two clone-mixed clumps (PA03 (*a*) and PA07 (*b*)) of *P. alexanderi*. Blue bars show the distribution in the first-instar normal nymphs and orange bars in the first-instar soldier nymphs. *P*-values of the dip test are shown.

## 4. Discussion

We report here the first detailed study of the clonal structure of social aphids living in open colonies and provide some insights into how eusociality might, uniquely within animals, have arisen in the absence of a nest. We studied two aphid species, which have colonies that feed on bamboos: the non-eusocial *Astegopteryx bambucifoliae* and the eusocial *Pseudoregma alexanderi*, which has sterile horned soldiers. We will first consider the clonal structure of the two species at three different spatial scales.

### 4.1. Clonal structure within aphid colonies

The clonal structure within the colonies of the two species was relatively similar. 89% (16/18) and 77% (10/13) of the colonies of *A. bambucifoliae* and *P. alexanderi* respectively consisted of multiple clones (two to 20 in *A. bambucifoliae*, two to six in *P. alexanderi*). Average pairwise relatedness within the colonies was 0.54 in *A. bambucifoliae* and 0.71 in *P. alexanderi* (Figure 2). Clonal mixing has been found in many open-colony-forming, non-social aphids (Loxdale and Brookes 1990; Guillemaud et al. 2011; Vantaux et al. 2011) and our results are consistent with this.

### 4.2 Clonal structure of aphid colonies on the same bamboo clump

There are interesting and significant differences between the clonal structures of the colonies of the two species at this spatial scale. The populations of the eusocial aphid *P. alexanderi* on large clumps of bamboo are much more genetically homogeneous than are those of the non-eusocial aphid *A. bambucifoliae*. In *P. alexanderi*, but not in *A. bambucifoliae*, the average pair-wise relatedness of aphids from the same small colony is not significantly different from the relatedness of two aphids from widely separate colonies on a large bamboo clump. In other words, a clump of bamboo typically contains several different clones of *A. bambucifoliae* but a single large colony of very restricted genetic diversity in *P. alexanderi*. Of the two species, therefore, an individual *P. alexanderi* roaming around a bamboo clump is much more likely to encounter a clone-mate.

### 4.3 Clonal structure across different bamboo clumps

There are significant differences between the two species at this larger spatial scale. In the non-eusocial *A. bambucifoliae*, each clone was restricted to a single bamboo clump, whereas in *P. alexanderi*, three of the four major clones were found at multiple sites (Table 2), which were more than 1km apart from each other.

### 4.4 Causes of the differences in clonal structure

The contrasting life-cycles and dispersal modes of the two species almost certainly underlie these differences in clonal structure (Figure 5). In *A. bambucifoliae*, colonies are founded by individual winged aphids flying from the primary host (*Styrax*) to the secondary host (bamboo) and it seems that there is little direct dispersal by aphids between separate bamboo plants. In Taiwan, the secondary host generations of *P. alexanderi* have lost all connection with the sexually-produced generations on the primary host and are maintained on the bamboo plants entirely by parthenogenesis. Younger instars of *P. alexanderi* can disperse by walking or floating on the air to found new colonies on distant bamboo plants (Aoki et al. 1981). However, the colonies of *P. alexanderi*, the eusocial species, tend to coalesce into a large colony on a clump of bamboo, which are both larger and more genetically homogeneous than the colonies on bamboo of the non-eusocial species, *A. bambucifoliae*, which are regularly restocked by genetically distinct aphids flying from the primary host.

**Figure 5.**
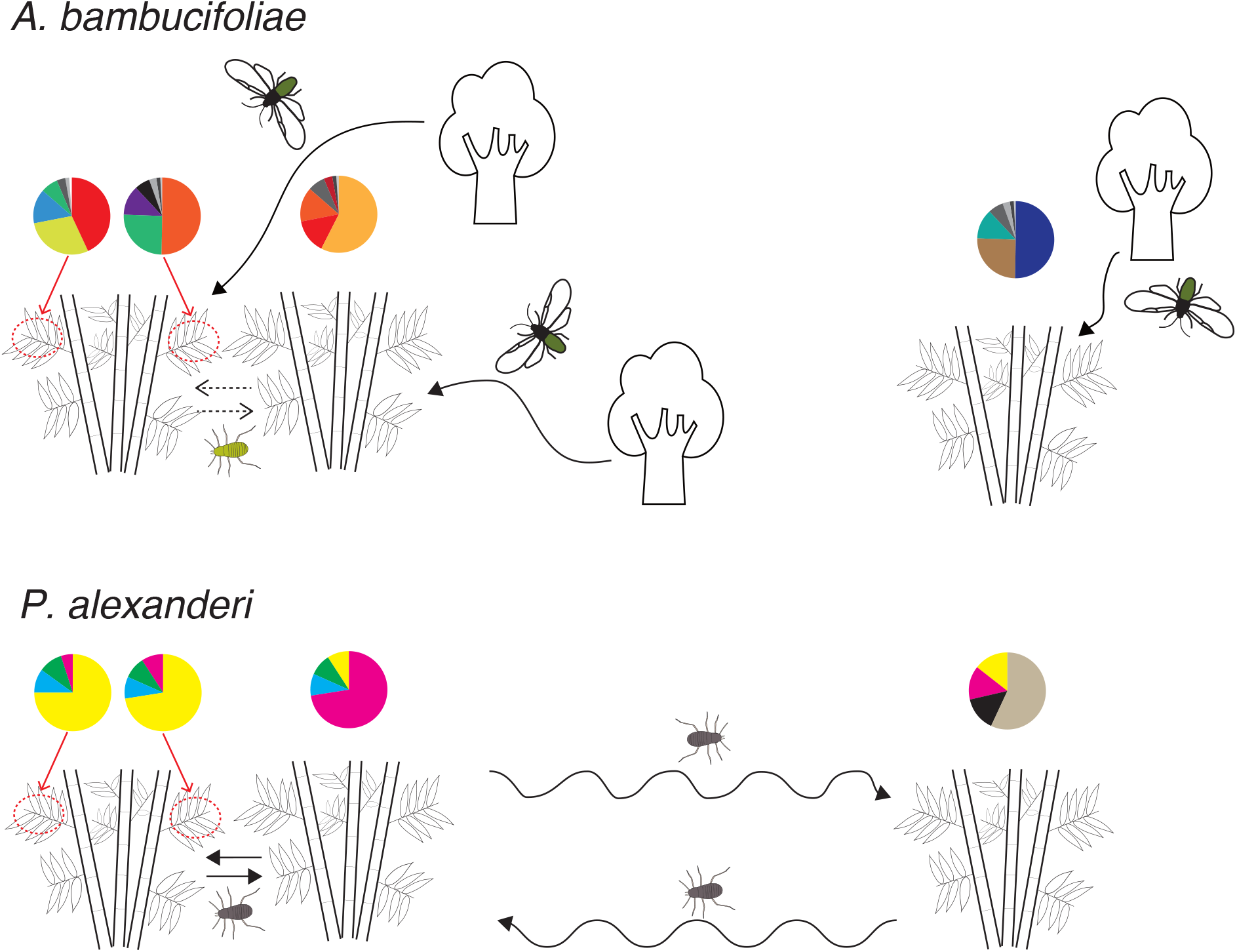
Comparison of the genetic structure between *A. bambucifoliae* (a) and *P. alexanderi* (b) within and between bamboo clumps. Colours in the pie charts indicate different clones. In *A. bambucifoliae*, winged adults frequently migrate from a distant gall-harbouring primary host plant, then mix with other resident clones in a bamboo clump. In *P. alexanderi*, the dispersing morphs are young nymphs that walk around between local clumps or migrate to long distance by walking or being borne by the wind. *P. alexanderi* has no primary host plant in this locality and can thus propagate only by parthenogenetic reproduction. As a result, colonies on the local site are genetically homogenised and major clones can spread to a distant bamboo clump in this species.

### 4.5 Consequences of differences in colony kin structure on soldier evolution in open colonies

Relatedness within a colony does not by itself determine the evolution of altruism, but it does set the threshold that the benefit to cost ratio of helping must exceed before altruism can evolve. Clonal mixing occurs in both species, but the average pair-wise relatedness within a colony of both species was higher than 0.5, the average relatedness between parent and offspring, and between siblings in a monogamous diploid family. These values, however, can be lower than relatedness values in colonies of some non-social aphids (0.90 in *Aphis fabae fabae*, 0.79 in *A. fabae cirsiiacanthoidis*) (Vantaux et al. 2011), although other studies that use different measures of genetic relatedness within a colony show higher degree of clonal mixing (Yao and Akimoto 2009; Loxdale et al. 2011).

Why has *P. alexanderi* opted to produce a sterile, morphologically specialized soldier caste, whereas *A. bambucifoliae* has not? Our genetic observations show that the major difference between the two species is that the eusocial *P. alexanderi* lives in populations that are relatively homogeneous genetically, even across different stems in a bamboo clump, whereas the non-eusocial *A. bambucifoliae* lives in colonies, each of which tends to be genetically distinct. A soldier aphid in *P. alexanderi* will therefore be much more likely to be defending a clone-mate, however far it has roamed on the bamboo clump, than is a defending first-instar of *A. bambucifoliae*. A defensive individual of *A. bambucifoliae* is quite likely to find itself defending a non-clone mate and the benefit/cost ratio for an average act of altruism will therefore be much lower than for a soldier of *P. alexanderi*.

### 4.6 Other ecological factors that might affect soldier evolution in open colonies

A second potentially important difference between the two species is simply that the colonies of *P. alexanderi* are much larger than the colonies of *A. bambucifoliae*. We regularly observed dense colonies of *P. alexanderi* on the bamboo stems, which from comparisons with related *Pseudoregma* species (Shibao 1998; Aoki et al. 2007) must have exceeded over 10,000 individuals. Some models suggest that, in any social organism, a soldier caste is more likely to evolve in larger colonies (e.g. Oster and Wilson (1978); Bourke, 1999; Ferguson-Gow *et al*.2014), but the evidence is mixed (Dornhaus, Powell & Bengston, 2012). In the social aphids, there is good theoretical and empirical support, from data on the development of individual colonies, for the idea that investment in soldiers increases as colonies grow larger (e.g. Aoki and Kurosu 2003, 2004; Aoki and Imai 2005; Sakata et al.1991; Schutze and Maschwitz 1991; Tanaka and Ito 1994; Shibao 1998). However, soldiers have been maintained in the open colonies of some Cerataphidini species with relatively small colonies (fewer than 1,000 aphids) such as *Ceratovacuna japonica* (Hattori et al. 2013) and *Pseudoregma panicola* (Aoki 1982). To understand soldier evolution in these relatively small colonies, future studies should investigate their genetic homogeneity on a large spatial scale. In addition, a phylogenetic comparative approach that included genetic homogeneity, colony sizes and other traits of many species of Cerataphidini could reveal whether a large colony favoured the initial evolution of eusociality.

A third factor that might be critical in the evolution of altruism in these open colonies is the nature of the host-plant and of the part of the plant upon which the aphids feed. Bamboos are long-lived perennial plants that grow and spread rapidly, producing a high biomass. Colonizing bamboo therefore enables long colony lifetimes and large colony sizes and it ensures the provision of vacant patches for dispersing aphids. In addition, it has been suggested that bamboo is a relatively poor food-source and that therefore investing in defensive behaviour is a wise strategy (Stern and Foster 1996), because the colonies are compelled to grow slowly. There is little direct evidence for this idea, but it is telling that the two species of horned aphids within the clade that have lost the soldier morph on the secondary host have both shifted from bamboo to herbaceous broad-leaved grasses (Stern 1998). This factor might be especially important for *P. alexanderi*, which grows on the hard woody stems and shoots of bamboo, rather than for *A. bambucifoliae*, which feeds on the relatively more nutritious leaves. This idea needs to be tested experimentally.

### 4.7 Maintenance of eusociality in open colonies

Social colonies that are not corralled within a nest face the additional problem of being especially vulnerable to invasion by free-rider unrelated conspecifics from another colony. This could be especially tricky for aphids which seem unable to distinguish clone mates from non-clone mates, in a variety of contexts (Aoki et al. 1991; Shibao 1999; Abbot et al. 2001). If this lack of kin recognition is genetically based, it might be a consequence of Crozier’s Paradox (Crozier 1986; Field et al. 2018). This posits that recognition cues will not evolve if individuals act as both hosts and invaders and if the costs of rejection are higher than the costs of acceptance, both of which might be true for these social aphids. There is some evidence that aphids are able to rely on the context in which they encounter another aphid. For example, first-instars of *Pemphigus* species appear to know when they are not in their home gall and behave appropriately (Abbot et al. 2001; Abbot 2009). An immature *P. alexanderi*, having been carried on the wind to a distant bamboo clump, might also “know” that it is no longer near its home clone, and behave accordingly. This is perhaps less likely in the non-eusocial *A. bambucifoliae*, which do not disperse over long distances between different bamboos and are more likely to encounter an alien colony after a relatively short and imperceptible excursion from their home clone.

Our results, admittedly from a small sample size, show that all of the four major clones of *P. alexanderi* contain soldiers in the mixed-clone clumps, suggesting that there is no obligate cheater clone that never produces soldiers. However, our results do not rule out the existence of conditional cheater clones that produce fewer altruistic soldiers when coexisting with other clones, as found in *Pemphigus* aphids (Abbot et al. 2001; Abbot 2009). In fact, the proportion of soldier and normal first-instar nymphs in the two mixed-clone clumps were significantly different among the clones, and the difference was consistent in both clumps, in which clone “A” produced fewer soldiers than expected, while clone “C” produced more (Table 4). This result suggests that there is genetic variation in soldier investment among clones, which could potentially adjust their investment of soldiers according to the presence of other clones in the same colony, although our observations provide no direct evidence for such an adjustment.

The existence of dimorphic soldiers (Figure 4) suggests that the open colonies of *Pseudoregma alexanderi* are relatively sophisticated for a eusocial aphid. Our results show that both the large and small soldiers are, like aphid morphs more generally, differential expressions of the same genotype (Figure S3). The functional significance of this dimorphism, which was not found in two related *Pseudoregma* species (Shingleton and Foster 2001), might be in providing useful phenotypic variation in the defensive effectiveness of the soldiers (Stern et al. 1996, Hattori et al. 2017). Interestingly, the most frequently observed clone “C” invested highly in the production of soldiers and is dimorphic. It would be interesting to see if these characteristics enhance the ecological success of a clone.

### 4.8 Future directions

The open colonies of social aphids provide an ideal model system in which to study the evolution of reproductive altruism. The critical parameters of the costs and benefits of soldier production and of relatedness can be measured and manipulated with relative ease. These parameters include predation pressure, colony growth-rate, colony size, soldier ratio and within-colony relatedness. It would be exciting to combine genetic analysis with manipulative experiments in *P. alexanderi* to determine the variability of the proportions of soldiers among clones and how this is influenced by the environment. There is a wonderful range of social aphids with open colonies, including the fairly distantly related *Colophina* species (Pemphigidae: Eriosomatinae) (Aoki 1977a,b; Uematsu et al. 2021). A comparative study of a range of these species on different host plants and with different degrees of sociality would greatly help us to understand the ecological and genetic factors that are important for the evolution of sociality in these unique and fascinating animals.

## Acknowledgements

This work was supported by supported by the Sumitomo Foundation (130972) and JSPS Overseas Research Fellowships (24-596) to KU. We thank Shigeyuki Aoki, Utako Kurosu, Chun-I Chiu, Bao-Cheng Lai, Yi-Chuan Lee, and Sheng-Feng Lin, and Chang-Ti Tang for help with fieldwork.

## Data Availability Statement

Additional data and details of statistical analyses are available in the Dryad Digital Repository (https://doi.org/10.5061/dryad.2547d7wtm).

