## Supplemental figures for "Eusocial evolution without a nest: kin structure of social aphids forming open colonies on bamboo"

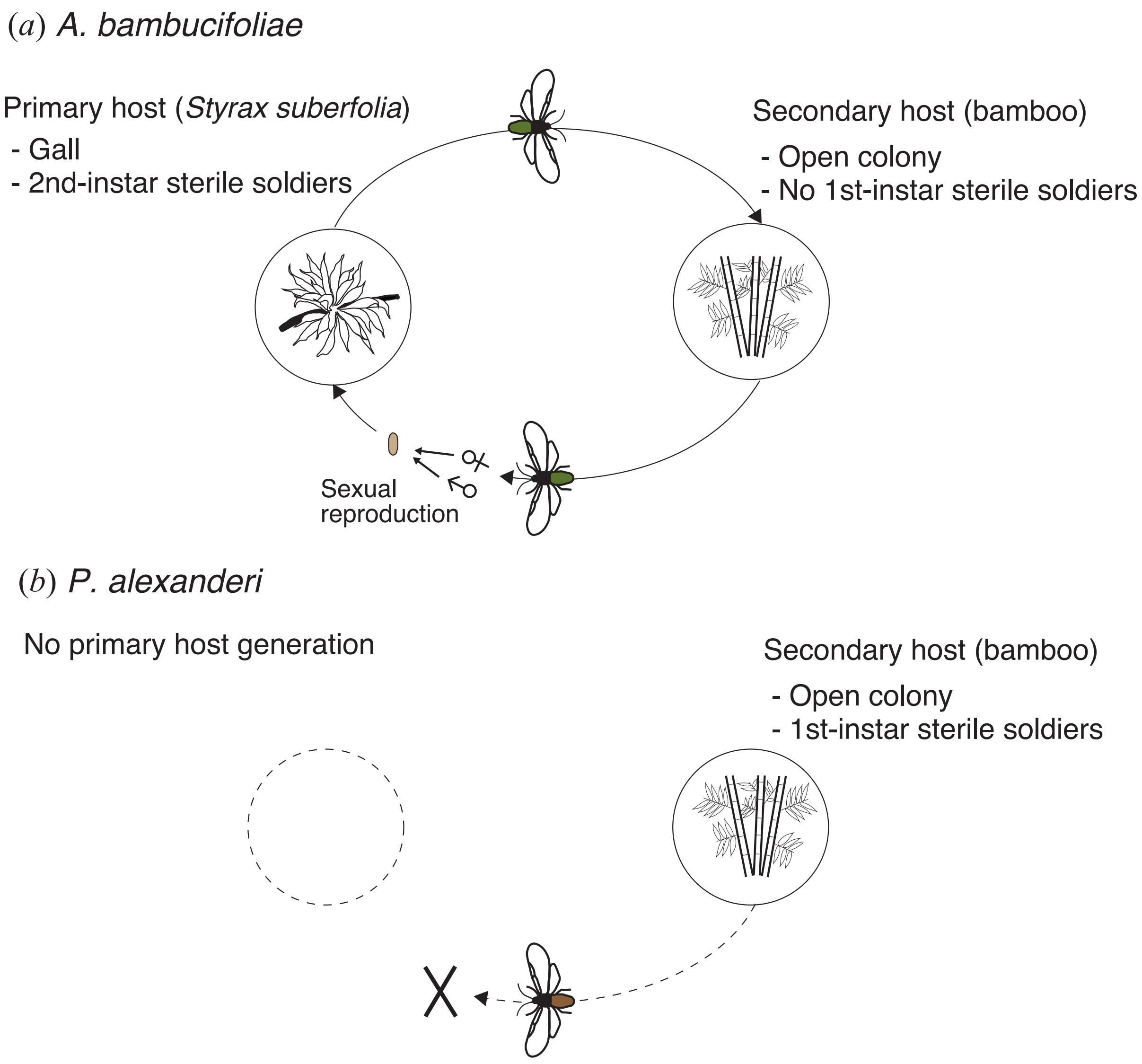


**Figure S1**. Life cycles of the two cerataphidine species in this study. Host-alternating life cycle of *A. bambucifoliae* (a) and non-host-alternating life cycle of *P. alexanderi* (b) are shown .

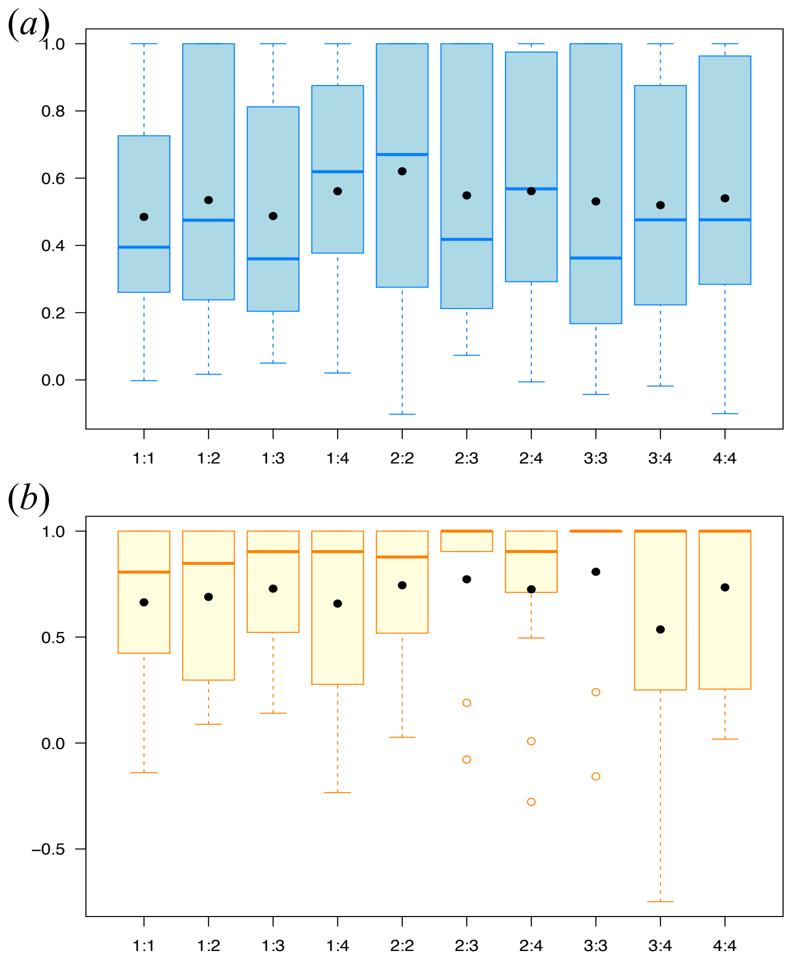


**Figure S2**. Boxplots showing pairwise relatedness values (*a*) for *A. bambucifoliae*, among first instars (1:1), between first instars and second or third instars (1:2), between first instars and third or fourth instars (1:3), between first instars and wingless adults (1:4), among second or third instars (2:2), between second or third instars and third or fourth instars (2:3), between second or third instars and wingless adults (2:4), among third or fourth instars (3:3), between third or fourth instars and wingless adults (3:4), and among wingless adults (4:4); (*b*) for *P. alexanderi*, among first-instar soldiers (1:1), between first-instar soldiers and first-instar normal nymphs (1:2), between first-instar soldiers and older nymphs (1:3), between first-instar soldiers and wingless adults (1:4), among first-instar normal nymphs (2:2), between first-instar normal nymphs and older nymphs (2:3), between first-instar normal nymphs and wingless adults (2:4), among older nymphs (3:3), between older nymphs and wingless adults (3:4), and among wingless adults (4:4). Boxes represent the interquartile range, bars within boxes are median values and whiskers indicate the 25th and 75th percentiles. Dots indicate mean value.

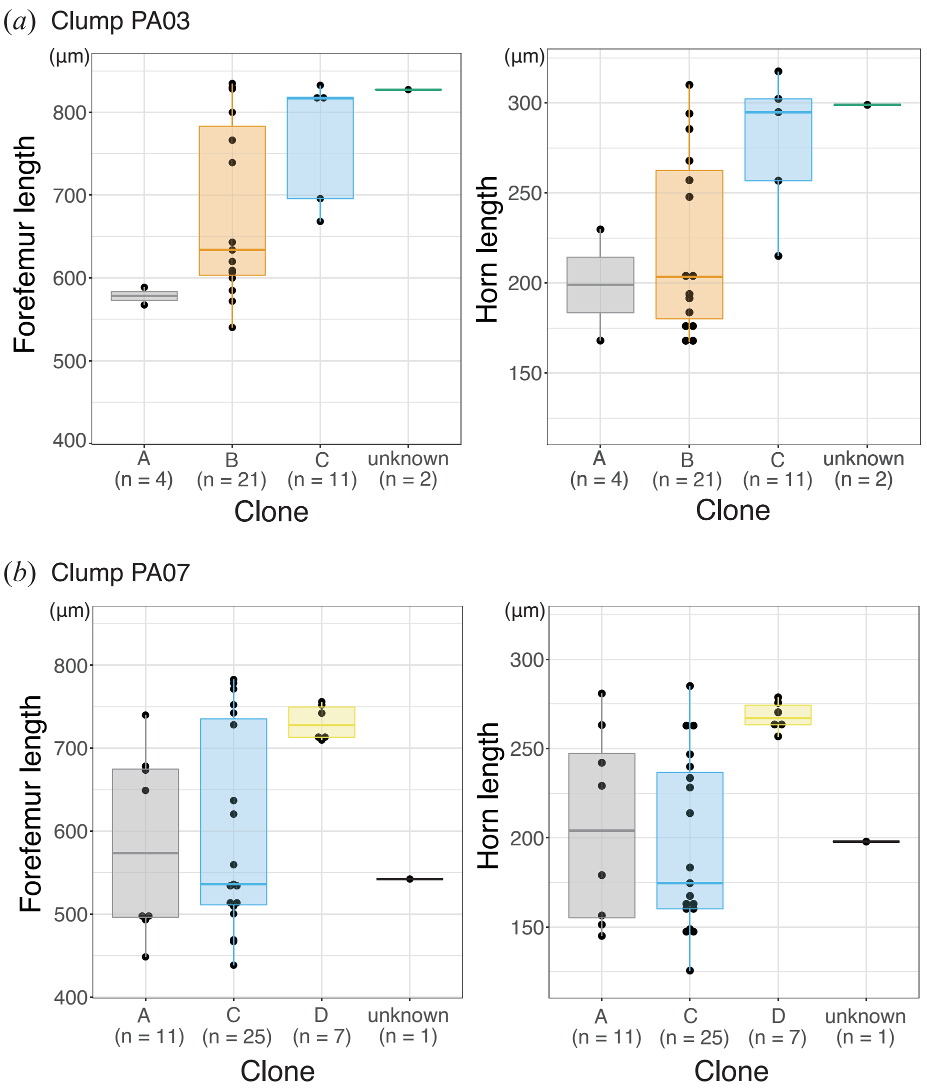


**Figure S3**. Box and dot plots of morphological characters of soldiers collected in the clump PA03 (*a*) and PA07 (*b*) and then genotyped. Fore-femur lengths (left) and horn lengths (right) of the genotyped soldiers of each clone are shown. The colors of the boxes represent clonal genotypes.
